# Clearing the way to the external world: do ants make optimal decisions when removing obstacles from their subterranean galleries?

**DOI:** 10.1101/2023.05.12.540516

**Authors:** Louis E. Devers, Clémentine Cléradin, Zoé Bescond-Michel, Gérard Latil, Vincent Fourcassié

## Abstract

Transport tasks are simple tasks whose cost can be easily measured and that are thus well suited to test optimality hypotheses. Here we focus on a particular type of transport that occur when ants are clearing obstacles from their subterranean galleries. In the laboratory we studied how they extract an object from a gallery of various inclinations. We expected that if ants behave optimally, they should remove the object by the gallery extremity requiring the lower energetic effort. At the colony-level, we found that the obstacle was more often extracted by the lower end of the tube, even if this required a higher amount of mechanical work. At the individual level however, ants showed mechanically optimal pulling behaviours in 75% of cases. Our results suggest that individual ants take into account both the inclination of the gallery and the position of the obstacle in it to decide in which direction they pull. In addition, they seem to base their decision to release the obstacle on the relative effort they perceive while pulling. Using a simple simulation model, we argue that the suboptimal extraction bias observed at the colony-level can be explained by the sequential nature of the extraction task.

## 1. Introduction

The optimality paradigm has been used for more than forty years in behavioral ecology as a powerful method to investigate how animals make decisions to solve a variety of problems involving cost-benefits trade-off, e.g. food or mate choice. However, the optimality approach has often been criticized for investigating too complex tasks and relying implicitly on unrealistic assumptions about the information gathering and cognitive capabilities of animals. Studies on insects are less prone to this criticism because, with their much smaller behavioral repertoire compared to vertebrates, the experimenter can focus on simple tasks that involve a limited amount of information. In addition, because of their small brain and limited lifetime, the experimenter is bound to make simple hypotheses on the cognitive mechanisms underlying the tasks to be optimized by these animals. Insects offer therefore a particularly good model for the study of optimality in behavior [1].

Social insects have been for a long time one of the favorite insects scientists have used for testing optimality theory. The optimality approach has been applied in these insects, both at the individual and collective level, to study decisions made in a variety of context such as reproduction, division of labor [2], food choice and foraging [3–6], colony defense, and nest choice [7] and construction [8]. One particular task that lends itself very well to the optimality approach is the transport of items. Ants, in particular, can transport all sorts of objects. They can transport food items such as whole prey, seeds or leaf fragments [9] but they can also transport building material [8,10], brood or nestmates [11] or various types of items when clearing their foraging trails of obstacles [12]. Each of these transports incurs a cost in terms of time and energy which is a function of the weight of the item transported and of the distance covered during the transport [13]. It also provides a benefit which can be assessed quantitatively by the energetic content of the item transported in the case of food or which is directly linked to the function of the object being transported in the case of a non-alimentary item.

In this paper, we focus on a particular type of transport that occurs in ants with subterranean nests when a narrow gallery linking their nest to the external world is partially clogged by an object, e.g. a small stone or a vegetation fragment, which has fallen down from the ground surface or which has been brought by the ants themselves. In that case the energetic cost required to transport the object in order to unclog the gallery depends on its inclination and on the distance to travel to reach either of its extremity. The benefit of the transport on the other hand lies in the reestablishment of the flow of ants in the gallery. In this situation, how do ants manage to extract the object from the gallery? do they behave optimally and extract the object in the direction associated with the less energetic cost? And on which basis do they decide to move the object in one or the other direction? In the study presented here we reproduced this situation in the laboratory and investigated the mechanisms by which ants, both at the individual and collective level, manage to extract an object from a narrow gallery with different inclinations. We used the measure of the mechanical work achieved while moving the object as a proxy for the measure of the metabolic energy expended. Since the displacement of the object can be done only in two directions, the relative cost of the two solutions can be easily calculated. We reasoned that if ants behave optimally when moving the object, they should choose, both at the individual and colony-level, the direction requiring the less amount of metabolic energy.

We structured our study in three different parts. First, we analyzed the final outcome of the sequence of individual pulling actions leading to the extraction of the obstacle to test whether ants reach energetically optimal solutions at the colony level. To do so we ran a series of experiments to calculate the position of the obstacle in the gallery for which the probability of left/right extraction was the same and compared it to the position for which equal amount of mechanical work was required to extract the obstacle from either end of the gallery. We then turned to the analysis of ant behavior at the individual level and examined whether the decision of individual ants to grasp and pull the object is also energetically optimal. Finally, we present a simple simulation model that aims at reproducing the clearing behavior observed at the collective level from the sequence of pulling actions observed at the individual level.

## 2. Methods

### (a) Studied species

The study was conducted on *Messor barbarus* (Linnaeus, 1767), a species of seed-harvesting ant commonly found in Mediterranean regions [14–16]. This species builds large subterranean nests and its colonies can be composed of several thousand individuals [15].

The worker caste of *M. barbarus* exhibits a continuous size polymorphism, with individuals ranging in length from 2 to 15mm and weighing from 1 to 40mg [17,18]. This size polymorphism can be roughly discretised in three size classes: *minor*, *media* and *major*. The *major* can be particularly well distinguished because of their much larger reddish head compared to the *media* and *minor*.

In total seven experimental colonies composed of 300 workers (150 *minor,* 100 *media* and 50 *major* individuals) drawn from five colonies collected in Saint-Hippolyte (France) between 2018 and 2020 were used in the experiments. The mean body mass of a sample of workers drawn from the experimental colonies was 21.40, 9.66 and 3.60 mg, for *major*, *media* and *minor* workers respectively.

### (b) Experimental set-up

Each experimental colony was installed in two transparent plastic boxes of identical size (H x W x L: 54 x 119 x 173mm) covered by a lid and linked together by a rigid tube (5mL Falcon® pipet, internal diameter: 6mm, length: 300mm). One box contained the nest and the other was used as a foraging area. Two plastic tubes were placed in each box. The two tubes in the nest box were used as nesting sites by ants. They were covered with opaque paper and had a water reservoir at their end retained by a cotton plug to provide water to the ants. The two tubes in the foraging box were entirely filled with pure water. In addition, ants had at their disposal a mixture of seeds and small cups filled with a mixed diet of vitamin-enriched food [19]. To avoid water condensation in the tube linking the two boxes, this latter was pierced at 18 mm interval with 2mm diameter holes that were covered with a fine metal mesh to prevent ants from escaping. Each box was placed on an adjustable platform so that the inclination of the tube between the two boxes could be set at a desired angle. In all experiments the nest box was placed on the left-hand side of the experimenter and the foraging box on the right-hand side. In addition, in the experiments in which the tube was not horizontal, the nest box was always at a lower position than the foraging box. Outside experimental sessions the experimental colonies were kept at a mean temperature of 22°C and relative humidity of 40% on a 12:12 L:D cycle.

### (c) Experimental protocol, data acquisition and data analyses

The experiments consisted in inserting an obstacle inside the tube linking the nest box and the foraging box and in waiting for the ants to remove it from the tube through one of its ends. The obstacle was a small piece of wood (diameter: 4mm, length: 60mm, mass: 300mg) that was inserted in the tube by one of its ends (either that of the nest box or that of the foraging box, chosen at random). The initial position of the obstacle and the inclination angle of the tube were varied between experiments. Each replicate of the experiment ended when the obstacle had been extracted from the tube by one of its ends or after one hour maximally. The connecting tube was changed for a new one at the end of each daily session of experiments to prevent the inner surface of the tube from too much wear due to the friction of the obstacle and of walking ants.

To find the point of equiprobability of left-right extraction of the obstacle and to investigate to what extent the position of this point depends on the initial position of the obstacle in the tube and of its inclination angle, the obstacle was inserted at five possible positions inside the tube, i.e., at 5, 10, 15, 20 or 25cm from the left (nest) end of the tube, and the inclination of the tube could take four different angles i.e., 0, 10, 20 or 30°. To determine the point of equiprobability of left(downward)/right(upward) extraction for each inclination angle of the tube at the colony level, we fitted a logistic regression with a logit link function. The response variable was the final destination of the obstacle in the experiments (set to 1 when the final destination was the right end of the tube, i.e. the foraging box, and to 0 when it was the left end, i.e. the nest box) and the inclination angle of the tube, the obstacle initial position and the interaction between these two variables were entered as independent variables in the model. We then used the equation of this model to compute, for each inclination angle, the position of the obstacle for which ants have equal probability to extract the obstacle to the right or to the left of the tube. We considered this position as the point of equiprobability of left/right extraction.

To minimize the error in the location of the point of equiprobability, we adopted a high throughput experimental design by working with four experimental colonies in parallel. For each colony a total of at least 200 replicates of the experiment were run with a given inclination angle and initial position of the obstacle. The inclination angle of the tube was chosen pseudo randomly so that at any time each colony was tested with a different angle. A total of 851 replicates of the experiments were performed on the four colonies. Among these 851 replicates, the obstacle was extracted from the tube in less than one hour in 792 replicates (93%).

To investigate how individual ants behave towards the obstacle as a function of its position in the tube and of the inclination angle of this latter, we used a different experimental design in which only one colony was tested at a time. First, we ran 144 replicates of the experiment in which the initial position of the obstacle was always set in the middle of the tube, which could be randomly positioned at four possible inclination angles (0°, 10°, 20°, 30°). Second, to create as much variability in the initial position of the obstacle, we changed its initial position systematically between each replicate, depending on the result of the preceding replicates. The result of each replicate was noted as one if the final destination of the obstacle was the foraging box and as zero if it was the nest box. Then, for each inclination angle, we fitted the results with a logistic regression and the equation of the logistic model gave us the theoretical position of the obstacle in the tube for which the odds of extracting the obstacle towards one or the other end of the tube are equal. This position was used in the next replicate as the new initial position of the obstacle. All replicates of the experiment run with this experimental design were videotaped at 50 fps using a JVC GZ-MG505 camera (resolution: 2560 x 1920 pixels) on three experimental colonies. Each experimental colony was tested at least six times before changing the inclination angle of the tube. A total of 120 replicates were run on the three experimental colonies with this method.

To acquire data on the behaviour of individual ants towards the obstacle we used the software Boris [20](https://www.boris.unito.it/). For each grasping event we noted the size class of the worker (*major* or *media* – detecting grasping by *minor* on the videos was too uncertain), its position relative to the obstacle (on its left or right side), as well as the time at which it began to grasp the obstacle and the time at which it released it.

To determine the position for which equal mechanical work was required to extract the obstacle from either end of the tube, we needed to calculate the forces applying on the obstacle. These forces depend on whether or not the obstacle is in movement.

When the obstacle is *not in movement*, it is subject to three physical forces (Fig. S1A):

- its weight *F*_*g*_ = *m* . *g*
- the reaction force of the tube *F*_*N*_ = *m* . *g* . cos(*α*)
- a static friction force *F*_*s*_ that prevents the obstacle from sliding along the tube when the tube is not horizontal

with *m* the mass of the object, *g* the gravity vector and *α* the inclination angle of the tube.

When an ant applies a force *F*_*A*_ on the obstacle, the static friction force increases in a direction opposed to the movement until reaching a maximum value *F*_*s max*_ just before the object starts to move (Fig. S1B). This force can be computed as the product of the coefficient of friction *μ*_*s*_ of the tube by its reaction force *F*_*N*_. Then, when the obstacle starts to move, one enters in a dynamical friction regime in which the coefficient of friction *μ*_*s*_ becomes slightly less important, thus decreasing the intensity of the friction force *F*_*s*_. For the sake of simplicity in our study, we estimated the friction force as the maximum static friction force (thus under the hypothesis that the object has a constant velocity).

To estimate the coefficient of friction *μ*_*s*_, we used the following method. We took each tube used in the experiments and we glued it to a vertical surface by plasticine at one of its ends while the other end was held by the experimenter to maintain the tube in a horizontal position. The obstacle was then inserted into the tube close to the attached extremity and the free extremity of the tube was released so that it slowly fell down. A camera was positioned in front of the tube to videotape the falling tube and determine precisely the inclination angle at which the obstacle started to slide down the tube. This angle, 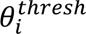 corresponds to the inclination angle for which the friction force applied on the obstacle are not high enough to prevent it from sliding. This operation was repeated 10 times for each tube and we estimated the coefficient of friction *μ*_*s*_ by the average value of the tangent of the inclination angles 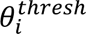. We then computed the maximal friction force exerted on the obstacle as **F*_smax_* = *μ*_*s*_ × *F*_*N*_.

Our data shows that the value of the coefficient of friction decreases with increasing exposure time of the tube to the ants (Fig S2B), ranging from 1.5 for a brand-new tube to 1 for a tube used for a total of 9 hours. The coefficient of friction in each replicate of the experiment was then estimated from the equation of a linear regression of the value of the friction coefficient against the duration of exposure of the tube to the ants.

From the estimation of the coefficient of friction *μ*_*s*_, one can then compute the minimal force *F_A min_* an ant has to apply on the obstacle to put it in movement for all inclination angles of the tube (0,10, 20, 30°) and the two pulling directions (upward or downward). When the tube is not horizontal, *F_A min_* is equal to the sum of the maximal friction force *F*_*s max*_ plus (if the obstacle is pulled upwards) or minus (if the obstacle is pulled downwards) the projection of the gravitational force on the tube axis *F*_*g*||_ = *m*. *g*. sin(*α*):

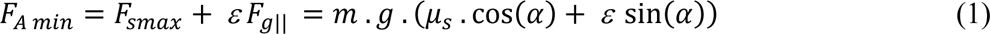

with ε= 1 if the obstacle is pulled upwards and ε= −1 if the obstacle is pulled downwards.

When the tube is horizontal, one gets α= 0 and thus sinα= 0. Consequently, the force that has to be applied on the obstacle to put it in movement is simply equal to the friction force.

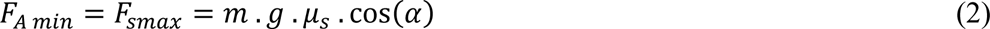

The mechanical work achieved by ants to extract the obstacle from the tube corresponds to the product of the force it has to apply on the obstacle by the distance it has to travel to reach the end of the tube towards which it is moving. The point of equal mechanical work for upward or downward extraction is thus given by the equation

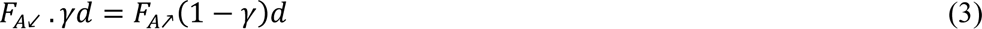

where *F*_*A*↙_ is the force that ants have to apply to extract the obstacle by the left (nest) end of the tube, *F*_*A*↗_ to extract it by its right (foraging box) end, and **γ*d* is the position of the obstacle in the tube (*γ* is the proportion of the length of the tube *d* = 30cm)

From equation (3) one gets:

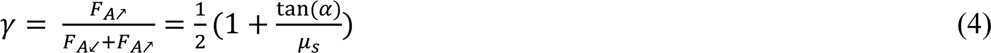

The position of the point of equal mechanical work can then be expressed as the distance 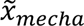 from the nest box:

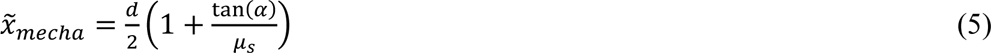

We performed all data analysis with R 4.2.1 (2022-06-23 ucrt) [21](RCore team, 2022) run under RStudio (v. 2022.07.0). The package *MuMIn* [22] was used to choose the most likely model among several regression models. In the presentation of the results the coefficients of the regression models are followed by the lower and upper value of the 95% confidence interval indicated in brackets. We used a Cox proportional hazard model to analyse the rate of release of the obstacle by ants. The survival analysis was run with the functions of the package *survival* [21] and the survival curves were drawn with the functions of the package *survminer* [23].

## 3. RESULTS

### (a) Colony-level extraction behaviour

Both the inclination angle of the tube and the initial position of the obstacle in it had an influence on the probability of observing an upward extraction in the experiments (Fig 1A, Table 1). When the obstacle was in the middle of the tube and the tube was horizontal, ants had equal chances of extracting it in either direction (*p*^^^ = 0.49, *N* = 41, *P* = 0.438). The downward extraction was increasingly preferred when the slope angle of the tube increased, even when the obstacle was placed initially close to the upward end of the tube.

**Figure 1.**
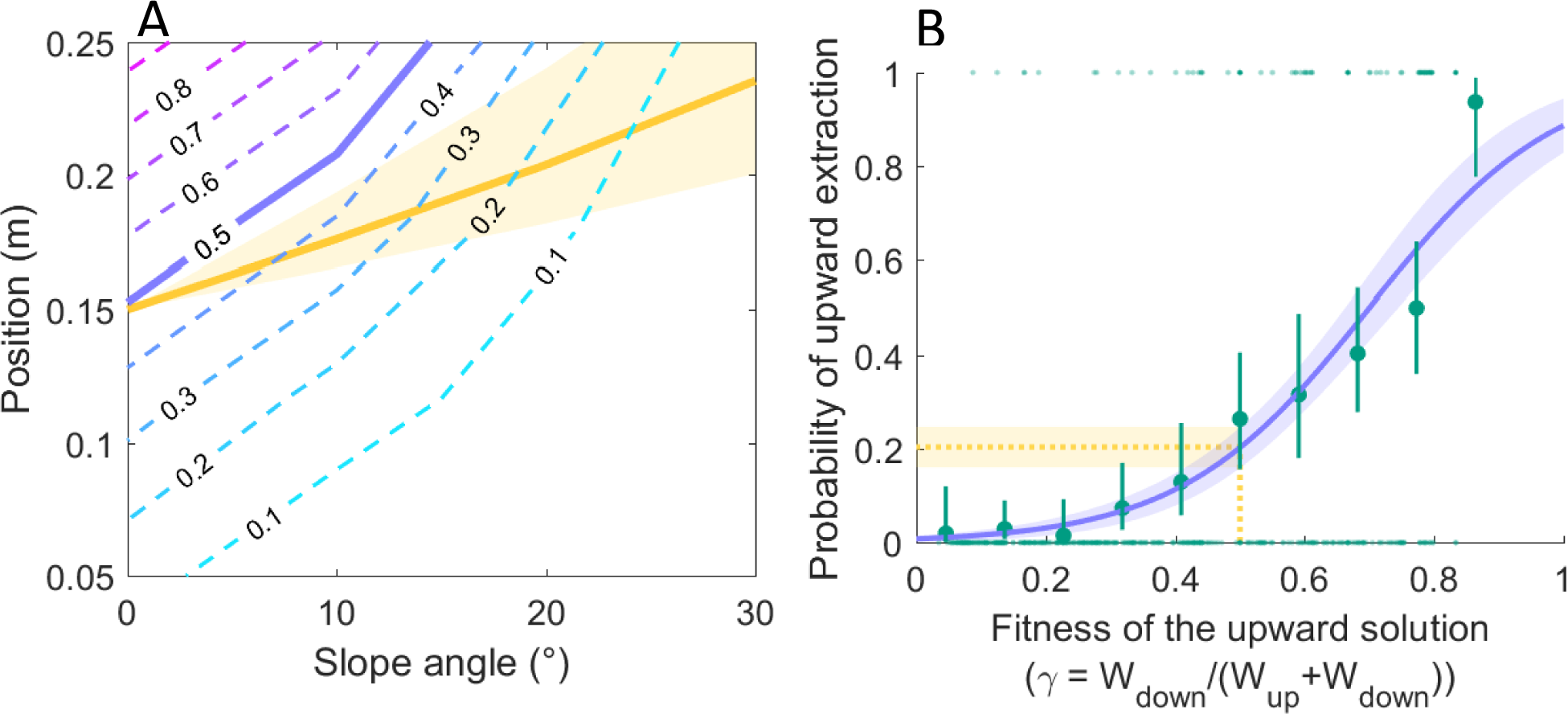
Colony-level extraction of the obstacle. (A) Probability of upward extraction of the obstacle as a function of both the inclination angle of the tube and the initial distance of the obstacle from the downward end of the tube (0m = downward end, 0.3m = upward end). The blue dashed lines are the lines of isoprobability of upward extraction. The solid purple line is the line of equiprobability of downward/upward extraction. The yellow line corresponds to the isowork line, i.e., to the values of slope and distance to the downward end of the tube for which the mechanical work required for an upward or downward extraction of the obstacle is the same. The yellow area around the line is the 95% confidence interval (based on the 95% confidence interval of the friction coefficient *μ*_*s*_) (B) Probability of upward extraction of the obstacle as a function of the mechanical fitness of the upward solution. The average empirical probabilities of upward extraction (along with their 95% confidence interval), calculated for 10 bins of 0.1, are shown as green dots. The purple line is the logistic fit surrounded by its 95% error bounds (purple area). The horizontal yellow dashed line shows the value of probability of upward extraction for which equal mechanical work is required to extract the obstacle by either end of the tube. The yellow area around the line is the 95% confidence interval (based on the 95% confidence interval of the friction coefficient *μ*_*s*_*). N* = 792 extractions.

**Table 1:**
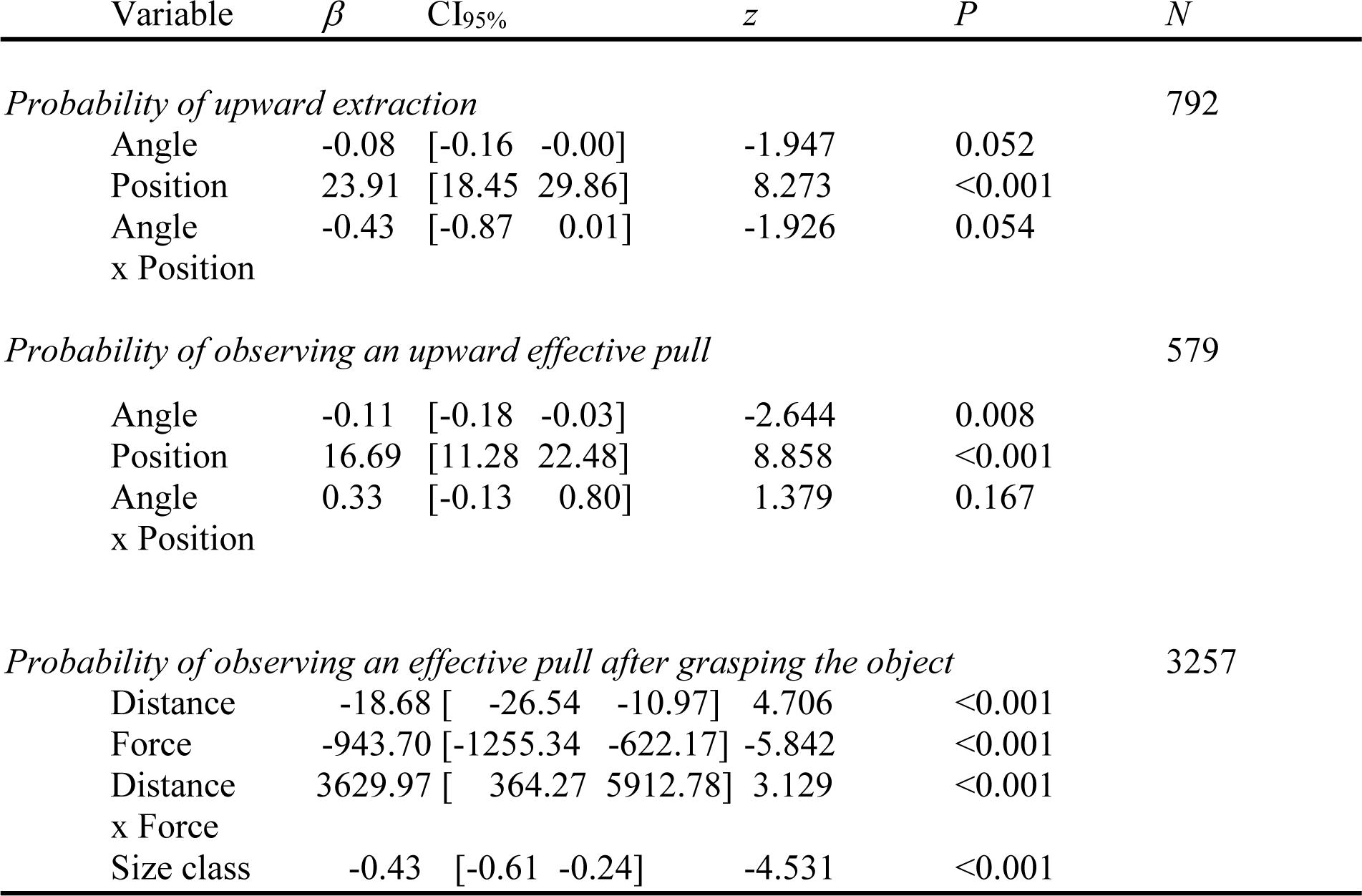
Output of model statistics for the regression analyses.

The comparison of the line of equiprobability of downward and upward extraction (Fig. 1A: solid purple line) with that for which equal mechanical work is required to extract the obstacle to either end of the tube (Fig. 1A: yellow line), shows that, except when the tube was horizontal, the direction of extraction was overwhelmingly suboptimal from a mechanical point of view, i.e. it did not correspond to the minimum mechanical work required to extract the obstacle. This suboptimal bias is shown in Fig 1B in which the probability of upward extraction is represented as a function of the mechanical fitness of this solution. The value of the probability of upward extraction corresponding to the point of equal mechanical work, i.e., 0.5 on the x-axis, is well below the value of equiprobable downward or upward extraction, i.e., 0.5 on the y-axis. At the colony-level, ants have thus a mechanically suboptimal extraction behaviour since they tend to extract the obstacle downward even when this requires more mechanical work than the upward extraction.

### (b) Individual pulling behaviour

All pulls did not necessarily lead to large displacement of the obstacle because ants sometimes released their grasp very rapidly. One can thus make a distinction between *aborted pulls*, that led to a very small displacement of the obstacle (≤ 1cm), and *effective pulls* that led to a larger displacement (> 1cm). In the remaining analysis only effective pulls will be considered.

Both the inclination angle of the tube and the position of the obstacle had an influence on the probability of observing an upward effective pull (Fig. 2A: Table 1). Upward pulling behaviours were adequately distributed from a mechanical point of view, i.e. ants did pull the obstacle upwards when the upward extraction required less mechanical work than the downward one. Besides, when both options require the same amount of work, ants responded randomly (Fig. 2A: yellow dashed lines and area).

**Figure 2.**
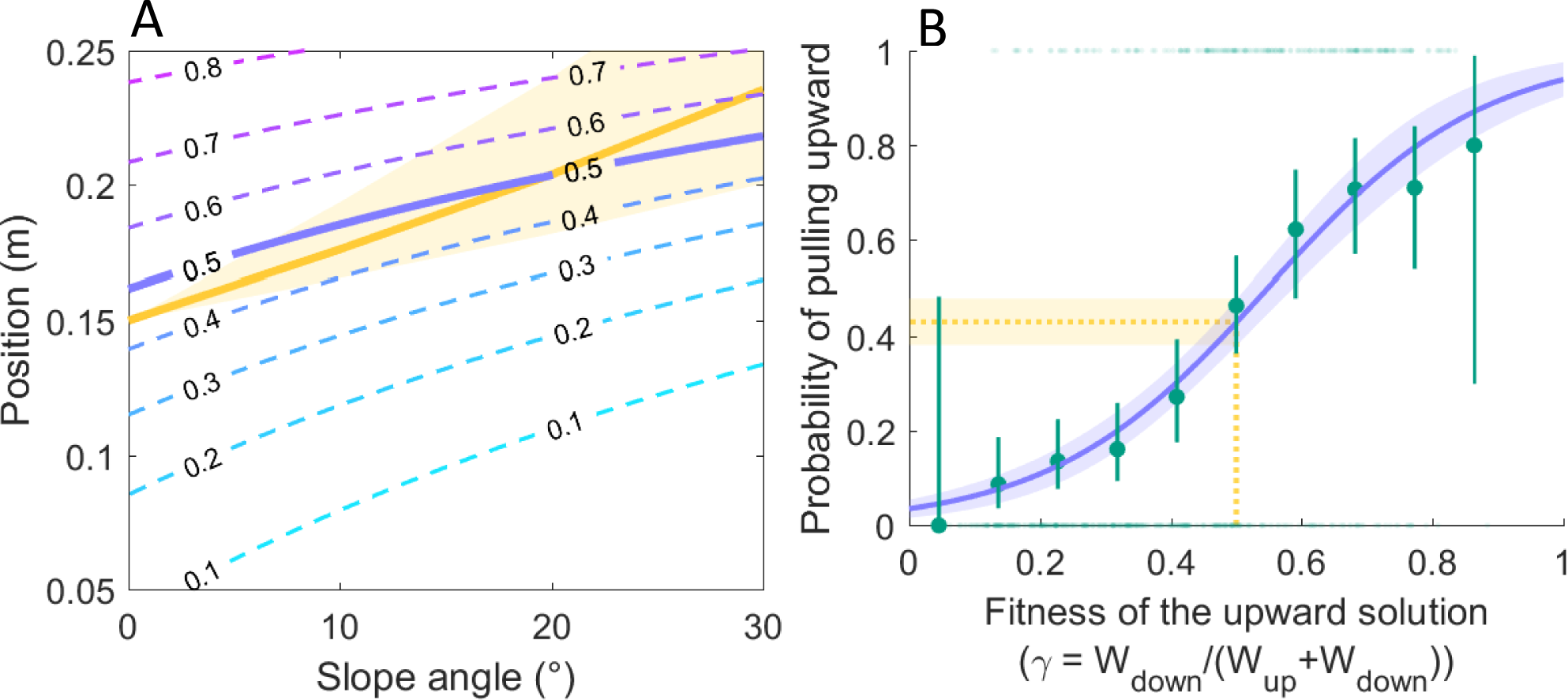
Individual pulling behavior. (A) Probability of observing an upward effective pull of the obstacle as a function of both the inclination angle of the tube and the distance of the obstacle from the downward end of the tube (0m = downward end, 0.3m = upward end). The dashed lines are the lines of isoprobability of upward pull. The solid purple line is the line of equiprobability of downward or upward pull. The yellow line corresponds to the isowork line, i.e., to the values of slope and distance to the downward end of the tube for which the mechanical work required for an upward or downward extraction of the obstacle is the same. The yellow area around the line is the 95% confidence interval (based on the 95% confidence interval of the friction coefficient *μ*_*s*_) (B) Probability of observing an upward effective pull on the obstacle as a function of the mechanical fitness of the upward solution. The purple line is the logistic fit surrounded by its 95% error bounds (purple area). The average empirical probabilities of upward extraction (along with their 95% confidence interval), calculated for 10 bins of 0.1, are shown as green dots. The purple line is the logistic fit surrounded by its 95% error bounds (purple area). The horizontal yellow dashed line shows the value of probability of upward effective pulls for which equal mechanical work is required to extract the obstacle by either end of the tube. *N*=579 pulls.

What kind of proxy could ants use to decide to continue to pull the obstacle once they have grasped it, i.e. to perform an effective pull? One could think of two types of information: the distance to the end of the tube towards which they are moving and the force they have to apply on the obstacle to set it in motion. We used a logistic regression to test to what extent these two criteria, considered independently or as a combination, could influence the probability of pulling the obstacle. The model thus includes as independent variables the force applied on the obstacle, the distance to the end of the tube and the interaction term between these two variables. We found that the probability to continue pulling depended on a combination of the force that has to be applied on the obstacle and of the distance to the end of the tube (Table 1, interaction force x distance). Therefore, ants have to somewhat integrate both metrics to take the decision to continue to pull the obstacle once they have grasped it.

Do ants of different sizes use the same decision criteria to decide to continue to pull the obstacle? We tested this by adding the worker size class (media *vs.* major) as a categorical variable in the logistic regression above. The results show that there was indeed an effect of worker size: all things being equal, major ants have a higher probability of continuing to pull the obstacle than media ants (Fig. 2B, Table 1).

Once ants have decided to pull the obstacle, they can decide to release it at any moment. One can hypothesize that the ants’ decision to release the obstacle could depend on the duration of the pull, on the distance travelled while pulling, or on the mechanical work achieved while pulling. Therefore, we tested three different survival models to explain the rate of release of the obstacle: in the first model we analyzed the rate of release of the obstacle as a function of the time elapsed since the initiation of the pull, in the second as a function of the distance moved while pulling and in the third as a function of both the distance moved and the force applied on the obstacle, equivalent to the mechanical work exerted. To investigate to what extent the rate of release could depend on worker size (major *vs.* media) and pulling direction (downwards *vs.* upwards - experiments with horizontal tubes were not considered in these analyses), these two variables were entered as independent variables in the three survival models.

Regarding the effect of worker size, we found that, independent of pulling directions, media ants held the obstacle on average for shorter durations (Fig. 3A, Table 2), shorter distances (Fig. 3B, Table 2), and for less amount of mechanical work (Fig. 3C, Table 2) than their major conspecifics. As for the effect of pulling direction, we found that, independent of worker size, when pulling downward (and thus applying a lower force than when pulling upward), ants tended to pull for shorter durations (Fig. 3A, Table 2) and on longer distances (Fig. 3A, Table 2) compared to when pulling upwards. However, ants released the obstacle after applying on average the same amount of mechanical work in the two different pulling directions (Fig. 3C, Table 2).

**Figure 3.**
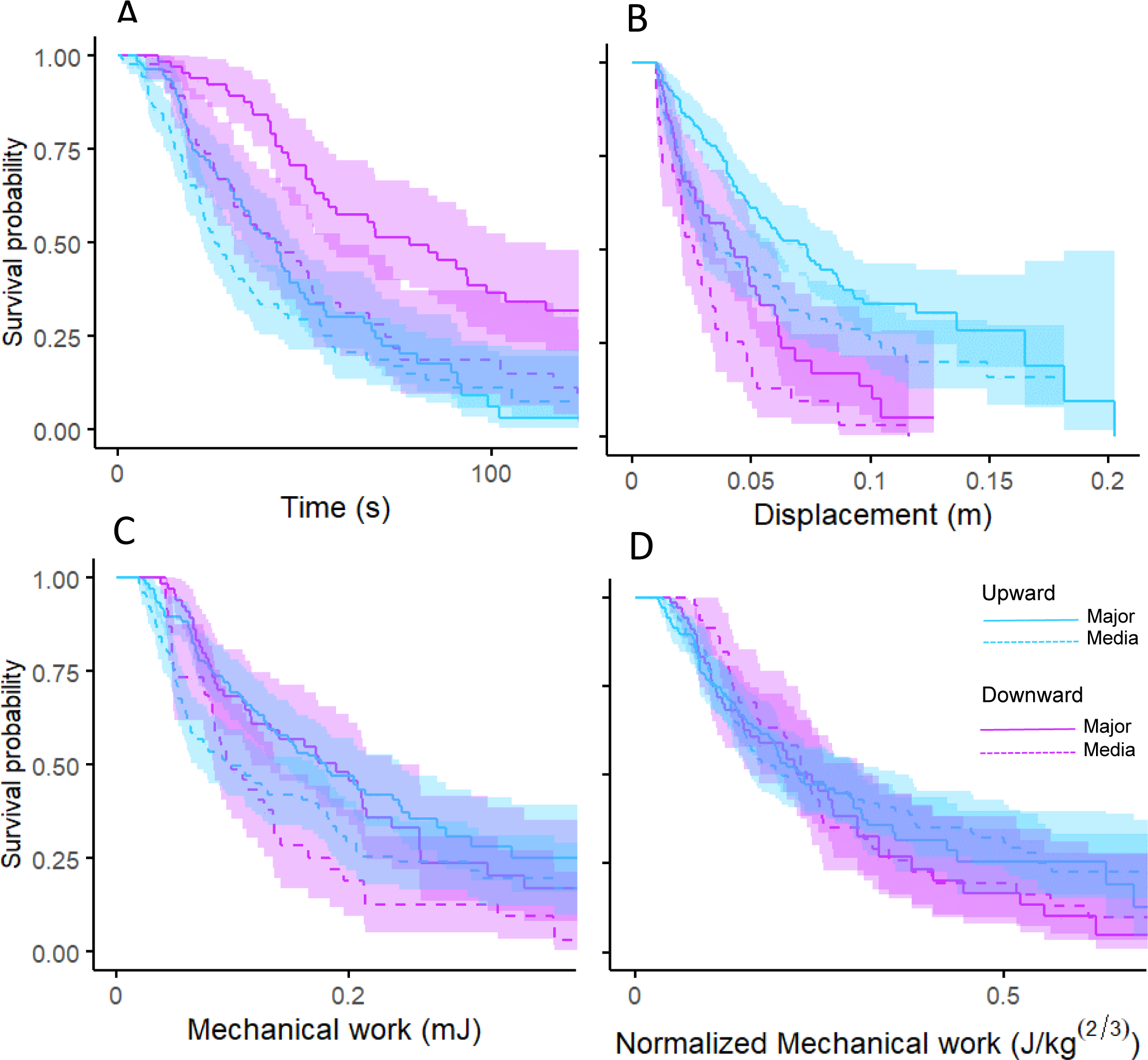
Kaplan-Meier survival curves of pulling behaviour, partitioned by worker size category (media/major) and pulling direction (upwards/downwards), as a function of (A) the time elapsed since the initiation of the pull, (B) the distance travelled while pulling and (C) the mechanical work achieved. (D) the mechanical work normalized by the mass^2/3^ of the workers. *N* = 729 pulls.

**Table 2:** Output of the Cox proportional hazard analyses on the rate of release of the object. The hazard ratio (HR) is the ratio of the rate of release of the object observed for media *vs.* major workers for the size class, and that of downwards *vs.* upwards pulling for the pulling direction. *N*= 729 pulls.

| Variable | HR | CI<sub>95%</sub> | z | P |
| --- | --- | --- | --- | --- |
| *Calculated on time* |  |  |  |  |
| Size class | 1.657 | [1.288 2.131] | 3.931 | <0.001 |
| Pulling direction | 0.486 | [0.370 0.638] | -5.190 | <0.001 |
| *Calculated on distance pulled* |  |  |  |  |
| Size class | 1.513 | [1.175 1.949] | 3.213 | <0.001 |
| Pulling direction | 1.916 | [1.443 2.544] | 4.498 | <0.001 |
| *Calculated on mechanical work* |  |  |  |  |
| Size class | 1.627 | [1.266 2.090] | 3.798 | <0.001 |
| Pulling direction | 0.875 | [0.670 1.142] | -0.985 | 0.324 |
| *Calculated on mechanical work normalized by body mass<sup>2/3</sup>* |  |  |  |  |
| Size class | 0.885 | [0.692 1.132] | -0.913 | 0.331 |
| Pulling direction | 0.856 | [0.673 1.089] | -1.263 | 0.207 |

### (c) Modeling colony-level extractions

Our study at the individual level shows that, on average, ants behave optimally from a mechanical point of view, i.e., most ants pull the obstacle in the direction requiring the less amount of mechanical work to extract it from the tube (Fig. 2A). In contrast, the extraction behavior at the colony level is mechanically suboptimal with a downward bias (Fig 1B). Therefore, the question arises of how a mechanically sound pulling behaviour at the individual level can lead to a suboptimal extraction behaviour at the colony-level. We argue that this is can be explained by the sequential nature of the task, i.e. to the repeated pulls and releases of the obstacle by many ants before it is eventually extracted from the tube. In fact, for each pull of the obstacle in the downward direction the probability for the next pull to be again in the downward direction increases more rapidly than if the pull was in the upward direction since, for the same amount of work, the obstacle is displaced on longer distance when it is pulled downwards than when it is pulled upwards.

To test whether this explanation is correct, we modelled the extraction behaviour as a sequence of successive downward or upward pulls over a certain distance until the obstacle is extracted from the tube. Each step of a sequence is a two-stage process. First, the decision to pull the obstacle is given by the equation of the logistic regression shown in Figure 2A, which gives the probability of observing an upward effective pull as a function of both the inclination angle of the tube and the position of the object in it. Second, the distance at which ants release the obstacle is based on the mechanical work they have achieved. Our model is based on observed data. Therefore, the mechanical work exerted by ants are randomly drawn from the distribution of the values of mechanical work observed in the experiment, independent of pulling direction and of the inclination angle of the tube. From the inclination angle of the tube and the pulling direction one can then calculate the force exerted by ants while pulling the obstacle and hence deduce from the mechanical work value the distance they travelled before releasing the obstacle. It is this distance that is used in the simulations to move the obstacle.

To compare the results of the simulations to those of the experiments, we reproduced in the simulations the same initial conditions as in the experiments (inclination angle of 0°, 10°, 20° or 30° and initial obstacle position ranging from 5 to 25 cm from the lower end of the tube). For each combination of angle and initial obstacle position the simulation was repeated 1000 times. We then compared the probability of upward extraction in our simulations for each combination of angle and initial obstacle position to that found in the experiments.

The results of the simulations are shown in Fig. 4. There was a relatively good fit of the model with the observed data, except for the highest inclination angle of 30°.

**Figure 4.**
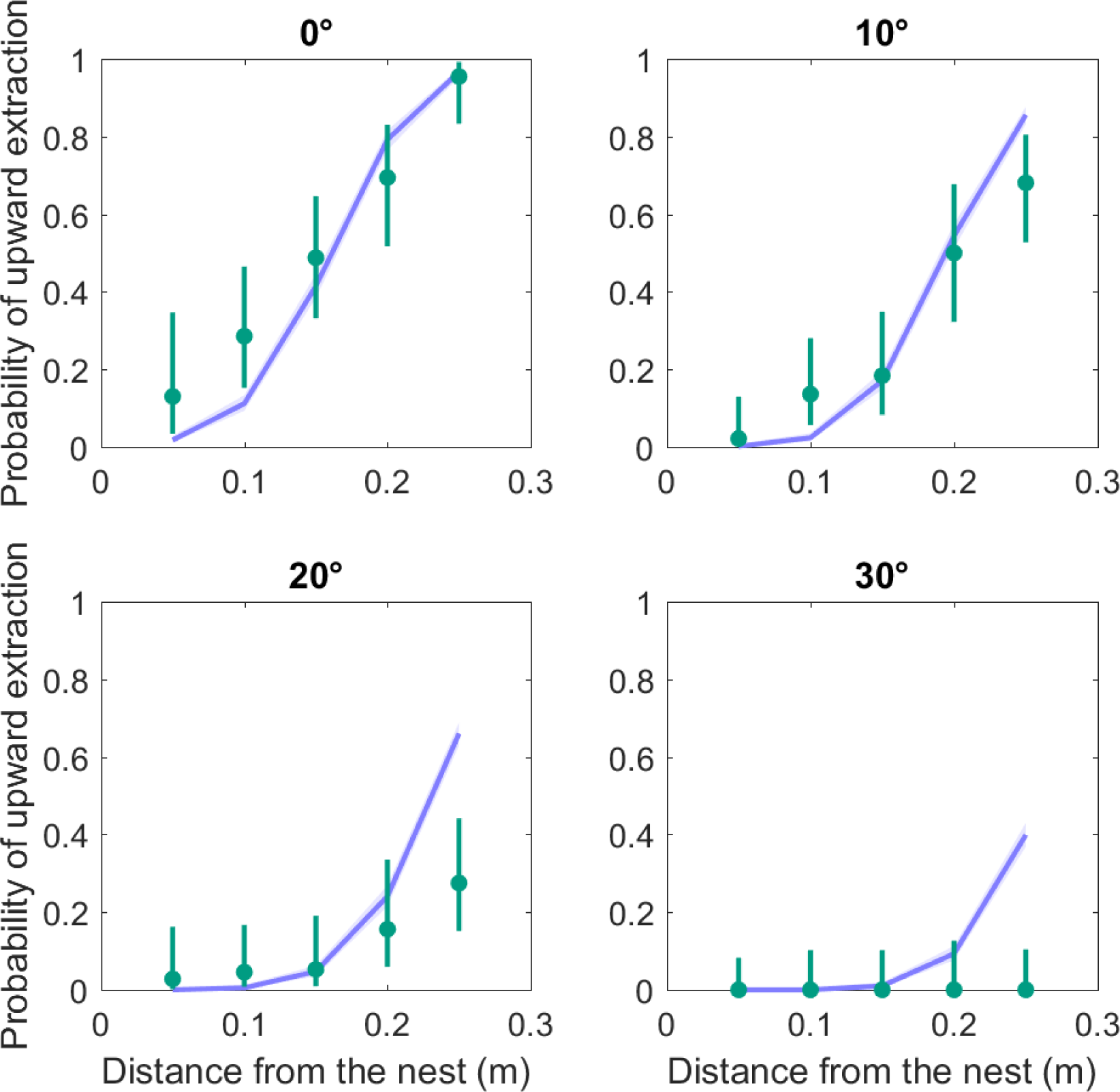
Comparison of the probabilities of upward extraction, calculated from the experiments and from the results of the simulations, as a function of the distance from the downward (nest) end of the tube and of its inclination angle. The green dots show the value of average empirical probabilities (along with their 95% confidence interval) calculated for five 5cm-distance bins. The probabilities calculated in the simulations are shown in purple. 1000 runs of the simulations were achieved for each combination of angle and initial obstacle position in the tube.

## 4. Discussion

When moving between their underground nest and their foraging ground many ant species use narrow galleries that can be temporarily clogged by all sorts of objects. These objects have to be rapidly removed to re-establish the flow of foraging workers between the nest and the ground surface. Here, we reproduced this situation in the laboratory and show that, at the individual level, ants behaved optimally by removing the obstacle most of the times in the direction requiring the less amount of mechanical work (Fig. 2), whereas at the colony-level a strong downward bias in the extraction of the obstacle was observed leading to mechanically non-optimal solutions (Fig. 1). Since ants were as likely to extract the obstacle by either end of the gallery when this latter was horizontal and the obstacle was positioned in its middle, this downward bias was not due to ants preferring to extract the obstacle in the nest direction but rather, as suggested by our simulation model, to the sequential nature of the extraction task.

How do individual ants make their decision to pull and release the obstacle in the gallery? Based on our analysis of the behaviour of individual ants, one can imagine a scenario in which each pull of the obstacle made by ants is the result of two successive decisions: i) the decision to grasp the obstacle and to pull it towards either extremity of the gallery, ii) the decision to release the obstacle at a new location after a certain time. Our analysis allows us to make the following assumptions regarding the criteria used by ants to make these two decisions. First, ants decide to continue to pull the obstacle once they have grasped it by considering *both* the distance they have to travel to reach the extremity of the gallery and the force they applied on the obstacle to set it in motion. This is actually equivalent to saying that they base their decision on an assessment of the total mechanical work needed to extract the obstacle from the gallery. Second, the result of our analysis of pulling durations allows us to make the assumption that, whatever the force they apply on the obstacle to set it in motion, ants base their decision to release the obstacle on the mechanical work applied since the beginning of the pulling action. Therefore, ants need to be able to assess two types of information to make mechanically optimal decisions: the position of the obstacle in the gallery and the force they have to apply on it to set it in motion. Below we discuss how these two types of information could be assessed.

There are two ways an ant could estimate the position of the obstacle in the gallery in our experiment. The first is by assessing its 3D location based on the surrounding visual cues that can be seen through the transparent wall of the tube used as a gallery. However, this hypothesis can be rejected because the visual landmarks present in the experimental room were too far away to allow for a precise assessment of the obstacle location in the tube by triangulation. The second mechanism which could be used by an ant to estimate the position of the obstacle is odometry, i.e. the assessment of the distance walked between the entrance of the gallery and the location in the gallery where the obstacle was encountered. Indeed, there are now firm evidences that ants can assess distance with a podometer that acts as a stride integrator [24,25], including when they walk on inclined surface as in our experiment [26,27]. Although most studies on ant odometry have been achieved over distances of several meters, distance estimation in ants can also be quite accurate on short distances [28], even in complete darkness [29]. Therefore, ants could use their podometer to assess the distance of the obstacle to each gallery extremities in our experiment.

Besides the position of the obstacle in the gallery, the second element ants have to take into account to assess the mechanical work required to remove the obstacle from the gallery is the force they have to apply on the obstacle to set it in motion. Insects possess various types of mechanoreceptors distributed over their legs to precisely monitor the position and movement of their limbs during locomotion and to adjust their walking pattern to the characteristics of the substrate on which they are moving (slope, uneven terrain, obstacle crossing…). Some of these mechanoreceptors like the *campaniform sensilla* are known to encode the forces exerted by the leg muscles as cuticular strains [30]. Although mostly studied in cockroaches and stick insects, this type of mechanoreceptors is likely to be also present in Hymenoptera. They could provide ants with proprioceptive feedback on the heightened activity of their leg muscles when they are dragging an object, allowing them to gauge the force they need to exert on it to set it in motion.

As mentioned before, independent of the slope of the gallery and of whether they were pulling downwards or upwards, the results of the survival analysis on pulling durations suggest that ants base their decision to release the obstacle on the amount of mechanical work achieved since the beginning of their pulling action. To explain this result, one could hypothesize that the limiting factor for continuing pulling is either the amount of energy that can be mobilized per unit time, i.e., the metabolic power, or, alternatively that ants have a limited amount of energy available to pull the obstacle and thus that they release their grip before using up this energy. Below we examine these two hypotheses.

The mechanical power required by ants for pulling the object is equal to the product of the force they apply on the object to set it in motion by the speed at which they move. This corresponds in our experiments to about 4.52μW [4.11 4.94] and 4.30μW [3.97 4.63] for media and major workers, respectively. Assuming that the work efficiency of ant leg muscles is about the same as that of the cockroach *Periplaneta americana*, i.e. around 1% [31], this amounts to a metabolic power of 452.0μW [411.0 494.0] and 430.0μW [397.0 463.0] for media and major workers, respectively. The specific metabolic power is thus 46790.89μW.g^-1^ [42546.58 51138.72] and 20093.46μW.g^-1^ [18551.40 21635.51] for media and major workers, respectively. In comparison, the specific metabolic power required for a worker of the seed-harvesting ant *Pogonomyrmex maricopa* to walk at a speed of 3cm.s^-1^ is about 10000.00μW.g^-1^ [32], which represents 21% and 50% of the specific metabolic power of media and major workers while pulling the object, respectively. Therefore, at least in the case of media workers, the metabolic energy cost per unit time may well be a limiting factor to continue pulling the obstacle.

The second hypothesis to explain why ants decide to release the obstacle is that they would have at their disposal a limited amount of energy available. But how much energy do ants expend when pulling the obstacle and what does this amount of energy actually represent? In our experiments ants released their grip after producing on average 0.124mJ [0.112 0.137] and 0.157mJ [0.144 0.171] an of mechanical work, for media and major ants respectively. Assuming, as above, that the work efficiency of ant leg muscle is about 1%, this would result in the use of 12.40mJ [11.20 13.70] and 15.70mJ [14.40 17.10] of metabolic energy for media and major ants respectively. Note that these values of metabolic cost are probably somewhat underestimated since they are not based on the total mechanical work achieved by ants, which includes the mechanical cost of locomotion, i.e. the mechanical work required to accelerate the center of the mass of the ants [33] as well as the internal energy they use to accelerate their limbs during locomotion, but just on the mechanical work required to pull the obstacle. Nonetheless, the metabolic cost of locomotion is likely to represent only a small fraction of the cost of pulling the obstacle. If one hypothesizes that the metabolic cost of locomotion in *M. barbarus* is about the same as that in *Pogonmyrmex maricopa*, i.e. around 129 J.kg^-1^.m^-1^ [32], a *M. barbarus* worker weighing 9.66mg, the average weight of a media worker in our experiment, could walk on average 9.95m [8.99 11.00] with the amount of metabolic energy required to pull the obstacle. And for a major worker weighing on average 21.4mg the distance covered would be on average 5.68m [5.20 6.18]. These figures are a few meters less than the average length of foraging trails observed in *M. barbarus* [34]. There are no data available in the literature on the maximal distance foraging workers of *M. barbarus* can travel per hour. As in most ant species, they can probably perform several round trips between their nest and a seed patch within the same hour and, what is more, with a load in their mandibles on their way back to the nest, which requires more energy than unloaded locomotion [35]. However, in *M. barbarus* [36] as in other ant species carrying external loads [37–40] the existence of transport chains, i.e. food transfer between workers along the foraging trails, is well documented. This means that loaded ants may not necessarily travel the whole length of the foraging trails with their load which may be a strategy to spare energy. Of particular interest is the fact that in *M. barbarus* the first workers in the chain tend to have a high loading ratio [36]. Therefore, although the amount of metabolic energy expended by ants while pulling the obstacle is not negligible, it may nonetheless be a factor that could contribute to limit the duration of pulling.

In conclusion, the distance at which ants release the obstacle may depend both on the metabolic power and on the amount of energy required to pull the obstacle. However, ants have no way to directly measure metabolic power or metabolic energy. Therefore, what are the criteria they could use to decide to release the obstacle? One of the criteria one could think of is the relative effort they perceive while pulling. This relative effort is linked to the relative force exerted by ants while pulling. The force exerted by a muscle depends on its cross-sectional area which increases with body mass raises to the 2/3 power [41]. Therefore, if the decision to release the obstacle depended on the relative effort perceived by ants one would expect that the differences between major and media found in the rate of release of the object calculated on the mechanical work achieved (Fig. 3C) would disappear if the mechanical work were normalized by body mass raises to the 2/3 power. This is actually what happens (Table 2, Fig. 3D). This suggests that ants could use the relative effort they perceive while pulling as a heuristic to decide when to release the obstacle. A similar heuristic has been shown in the ant *Pheidole pallidula* whose scout workers modulate their trail-laying behavior to recruit nestmates as a function of the tractive resistance of the prey they find [42]. In the same way, it is likely that the size-matching between body mass and load mass observed in several ant species transporting external loads could be explained on the basis of an assessment of the relative effort. Further experiments would be needed to test this hypothesis. For example, one could think of manipulating the force required to pull the obstacle by using a rough substrate to increase the coefficient of friction of the tube or by attaching a thread to its two ends so that the experimenter can decrease or increase the force ants have to apply to set the obstacle in motion and pull it in either direction. One could also place individual ants in a respirometric chamber and use flow-through respirometry to measure gas exchanges during pulling to assess the metabolic cost of pulling.

Our experiments show that the complete extraction of the obstacle from the tube by a single ant was very rarely observed. In fact, it was overwhelmingly the result of a series of successive pulls made by individual ants and it is the sequential nature of the task that explains the downward bias observed at the colony-level. The simulations we ran which modelled the extraction behaviour as a sequence of successive downward or upward pulls confirms this explanation. The poorest fit was obtained when the slope of the tube was 20° or 30° and the obstacle was very close to the upward end of the tube. The following hypothesis may explain this poor fit. In our model the decision to pull the obstacle at each step of a sequence is based on the probability of ants to perform an upward *effective* pull, i.e. to continue pulling once they have grasped the obstacle and started to pull it. However, when the force required to move the obstacle was maximum, i.e. when pulling upwards for the highest value of the inclination angle of the gallery, it is likely that many ants, especially media ants which have less force, quickly released the obstacle just after grasping it. The capacity of ants to pull the obstacle upwards for high inclination angles was thus overestimated in our model, leading to an overestimation of the probability of upward extraction.

In conclusion our study on gallery clearing shows that a mechanically optimal response at the individual level can sometimes lead through an amplification process to a mechanically non-optimal response at the collective level. This is reminiscent of the maladaptative response resulting from negative information cascade that are sometimes observed in group-living animals and that occur through direct [43] or indirect [44] information transfer. However, the situation we studied here is singular. While each individual makes a correct decision based on the relative effort required to move the obstacle, the maladaptive response at the colony-level occurs as a consequence of its action which is strictly governed by the law of physics.

## Data accessibility

## Author’s contributions

L.D. conceived the project and designed the experiments, L.D., and C.C. performed the experiments, L.D. and ZBM processed the videos, L.D. analyzed the data. V.F. and L.D. wrote the manuscript. V.F. and G.L. supervised the work.

