## Supplemental Figures S1&2 for "Clearing the way to the external world: do ants make optimal decisions when removing obstacles from their subterranean galleries?"

### SUPPLEMENTARY MATERIAL

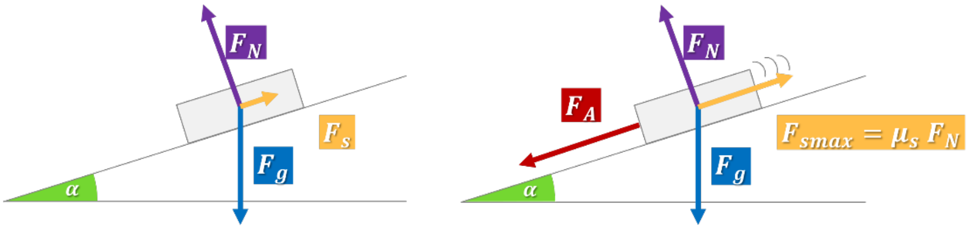


Figure S1 – Free body diagram showing the forces exerted on the stick when (A) the stick is at rest in the tube and when (B) an ant pulls it downward at a constant speed (null acceleration). α is the inclination angle of the tube, Fg is the weight of the stick, FN is the reaction force from the tube, FS is the static friction force and FSmax is the static friction force at its maximum value, just before the stick is set in movement.

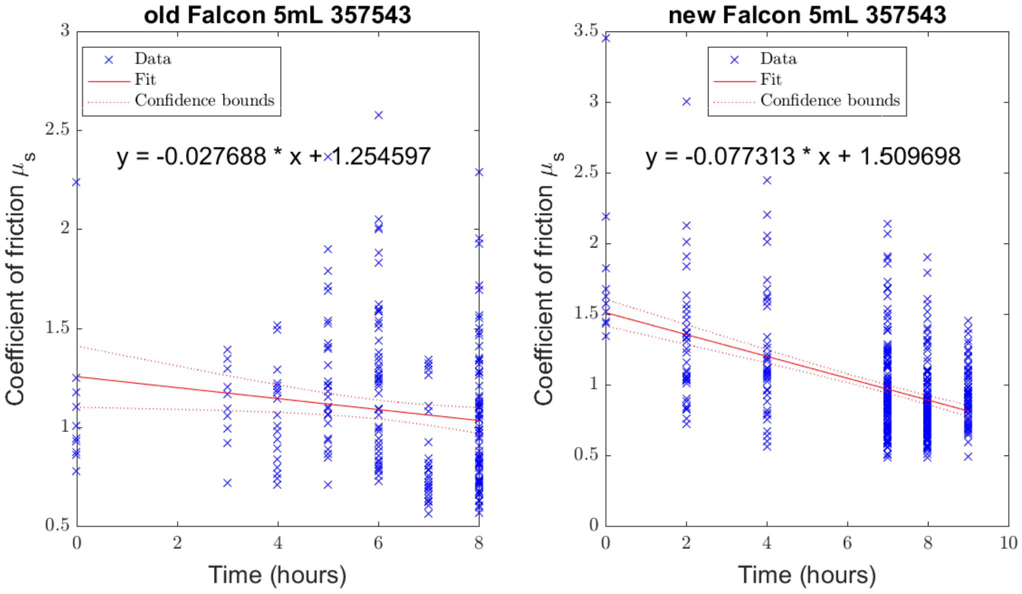


Figure S2 – Determination of the coefficient of friction of the tube $\mu_s$ as a function of the number of hours the tube has been exposed to ants in the experiment. The red line shows the predictions of a linear regression. Regressions were made for respectively old (A) and new (B) Falcon tubes of reference 357543.
